# Metagenomic Profiling of Municipal Drinking Water Microbiomes in an Indian City: Insights into Diversity, Water Quality, and AMR Potential

**DOI:** 10.1101/2025.02.03.636216

**Authors:** Vikas Kumar, Shradha Sharma, Kaomud Tyagi, Barathi Lenin, Aarti Ravindran, Karthik Raman, Inderjeet Tyagi

## Abstract

Monitoring microbial components in drinking water is as essential as tracking its chemical composition. Although traditional culture-based methods provide valuable insight into microbial morphology and behaviour, their scope is restricted to culturable species. With the advent of high-throughput sequencing, we can now detect a wider range of microbes in any ecosystem, along with efficient insights into their functional potential and metabolic capabilities. In this study, metagenomic analyses were performed to fully understand the microbiome of drinking water supplied through public distribution systems in an Indian city. Our findings identified bacteria from the phyla Pseudomonadota, Planctomycetota, Bacteroidota, and Actinomycetota, consistent with previous studies of drinking water microbiomes of other countries. At the species level, *Afipia carboxidovorans, Klebsiella pneumoniae, Pseudomonas aeruginosa, Sphingopyxis macrogoltabida*, and *Variovorax paradoxus* were identified as members of the core microbiome. It was observed that the temperature of the water samples, even as little as a 5ºC increase, influenced the composition and diversity of the microbial communities. No significant correlation was detected between the abundance of microbial species and the metal concentration in the sample. In addition, we traced the distribution of antibiotic resistance genes (ARGs), finding widespread resistance to aminoglycosides, tetracyclines, and macrolides in samples. In particular, ARGs such as adeF and ermR, which are known to be associated with multidrug resistance, were detected. Although this study did not directly assess the pathogenicity or mobility of these genes, their presence in potable water raises potential public health concerns due to the possibility of horizontal gene transfer (HGT) in environmental settings. Therefore, continuous monitoring of antibiotic resistance genes (ARGs) is imperative to accurately evaluate long-term risks and to guide evidence-based water quality management strategies. In summary, this study provides a comprehensive metagenomic overview of drinking water microbiota, ARGs, and water quality, offering a foundation for future surveillance and risk mitigation strategies.

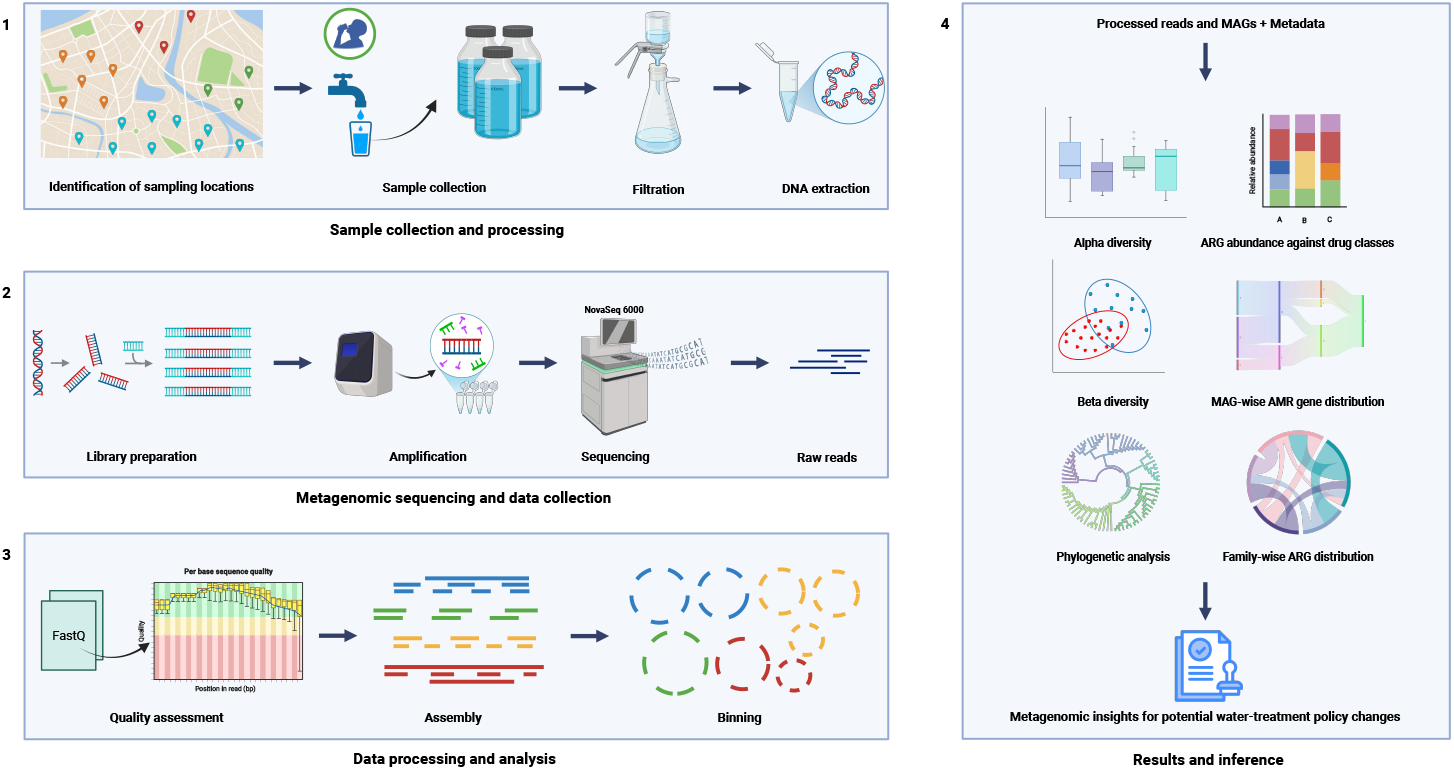

## 1 Introduction

Providing safe drinking water to all is a fundamental objective of public health agencies across the world, as it supports population health, socio-economic development, and prevents waterborne diseases. The global monitoring of drinking water, sanitation, and hygiene, collectively known as WASH, is a critical focal point of the World Health Organization (WHO)[1]. Ensuring the availability of safe and clean drinking water is a crucial priority, especially for a densely populated country such as India.

Robust drinking water treatment processes (DWTP) are employed to effectively treat water so that it becomes fit for consumption. The design of DWTPs involves managing the physical, chemical, and biological contaminants in untreated water. Briefly, physical processes such as sedimentation and filtration facilitate the removal of particulate matter. Chemical treatments eliminate pollutants such as dissolved harmful substances like Polycyclic Aromatic Hydrocarbons (PAHs), Per- and polyfluoroalkyl substances (PFAs), heavy metals like lead (Pb) and mercury (Hg), and synthetic cationic and anionic dyes. Biological contaminants include organic debris and a variety of organisms, typically bacteria, viruses, protozoans, fungi, and algae [2]. Methods such as adsorption, disinfection, filtration, flocculation, chemical treatment, and UV irradiation are commonly used to effectively remove pathogenic microorganisms that cause waterborne diseases [3]. However, the microbiological component of water consists of more than just live organisms, making it essential to address and neutralize associated elements that adversely impact human health.

Both free-living and biofilm-forming microbes can contain ARGs. Moreover, HGT plays a critical role in disseminating anti microbial resistance (AMRs) in the drinking water microbiome [4]. HGT can occur through mechanisms like transformation, transduction, and conjugation, leading to the spread of ARGs even in the absence of live pathogens. Additionally, the microbial load also comprises spores, endotoxins, exotoxins, mycotoxins, and phycotoxins - metabolites produced by certain microorganisms and can contribute to long-term health risks if ingested [5].

Addressing the microbial safety of water also aligns with the Sustainable Development Goals (SDGs) adopted by the United Nations in 2015. Specifically, SDG 3 was created to promote good health and wellbeing, while, SDG 6 was formulated to make water universally accessible and sustainable, both of which are directly impacted by water quality. Achieving these goals requires not only robust infrastructure, but also continuous monitoring and effective implementation of drinking water treatment and supply systems [6, 7].

In India, the Bureau of Indian Standards (BIS) regulates the quality of drinking water, with the latest amendment to the standards made in 2015 [8, 9]. Despite substantial improvements in water infrastructure, such as those driven by the Swachh Bharat Mission and the Jal Jeevan Mission (Ministry of Jal Shakti), there are still gaps in understanding, especially the microbial components of treated drinking water [8, 9].

In particular, metagenomic surveillance and analysis have not been performed to monitor the quality of drinking water in India. Our study is the first to carry out metagenomic sequencing of samples collected from drinking water outlets in a metropolitan city to understand the microbial component of treated water. This approach will enhance our understanding of potential risks and provide valuable insights to guide future improvements in water safety monitoring.

## 2 Materials and Methods

### 2.1 Collection of water samples and DNA extraction

Drinking water samples (municipal tap water) of approximately 10 liters each were collected in triplicate from 21 locations (n=63) selected based on a grid map approach in a metropolitan city located in India during March 2022. The exact locations are not disclosed. The samples were collected in high-density polyethylene (HDPE) autoclaved sterile bottles marked W1-W21 based on the collection sites. They were maintained at 4ºC and transported to the laboratory within three hours. The samples were then filtered immediately using 0.22 µm MCE membrane (Merck Life Science Pvt. Ltd., Bangalore) with the help of vacuum pump, and the biomass-containing membranes were placed into DNA extraction tubes and stored at -20 ºC till DNA extraction. Extraction of DNA from the samples was performed using the DNeasy PowerWater DNA isolation kit (Qiagen Cat No.: 14900-100-NF) following the standard manufacturer protocol, and the triplicates were pooled into 21 drinking water samples for sequencing. Pooling was performed due to the limited resources available for this study. The quality and quantity of extracted DNA were evaluated through agarose gel electrophoresis (Cell BioScience Alphalmager MINI) and a Qubit® Fluorometer (Invitrogen, USA), as described in previous studies [10]. In addition to the drinking water samples, a negative control was processed following the same protocol. However, it was excluded from sequencing as amplifiable DNA was not obtained for sequencing from this sample.

### 2.2 Library preparation, Whole genome sequencing (WGS), and preprocessing of sequencing reads

Using a Covaris ultrasonic disruptor, 1 µg of genomic DNA was randomly fragmented into segments of approximately 350 bp to construct the library. Subsequent steps in library construction were end repair, A-tail addition, sequencing adapter ligation, purification, and PCR amplification. Following this, AATI analysis was used to evaluate the size and integrity of the inserts and the library as a whole. Library quality assessment was performed using qPCR, which determined an effective concentration of greater than 3nM, thereby meeting the requirements for downstream sequencing. Once the library passed the quality check, it was combined with other libraries on the basis of their effective concentrations and data output targets. Paired-end 150 bp (PE150) sequencing was performed on the Illumina NovaSeq 6000 platform. The obtained raw data were preprocessed using fastp [11]. Adapters were trimmed, and the paired-end reads were discarded in the following cases: 1) when there were more than 10% unknown nucleotides in each read; 2) when there were more than 50% low-quality nucleotides (base quality less than 5). The final set of clean sequences was submitted to NCBI under the BioProject ID: PRJNA1008921 with accession numbers SRR25754722 to SRR25754742.

### 2.3 Metagenomic assembly, binning, and quality control

To remove human contamination from the adapter-trimmed reads, BMTagger v3.101 aligned the reads with the hg38 reference genome. Read pairs in which only one read aligned with the human genome were excluded. Quality checks were performed before and after filtering using FASTQC v0.12.1 [12]. The filtered reads were assembled using the MEGAHIT v1.1.1 [13] assembler within the MetaWRAP pipeline [14]. MEGAHIT employs a multiple k-mer-based strategy, where smaller k-mers fill gaps in low-coverage regions while larger k-mers resolve repeat sequences, thereby enhancing the overall assembly accuracy. The default k-mers of lengths 21, 29, 39, 59, 79, 99, 119, and 141 were used. Assembly was performed per sample, and contigs shorter than 1,000 bp were removed from the analysis. The assembled contigs were then binned using three algorithms: CONCOCT v0.4.0 [15], MaxBin2 v2.2.4 [16], and MetaBAT2 v2.9.1 [17], each run independently with default parameters. These algorithms leverage tetranucleotide frequencies, coverage information, and sequence composition to achieve precise binning. The consensus approach integrates the strengths of each algorithm, maximising the capture of genomic diversity while minimising potential errors that could arise from using a single method. Each of these algorithms produced separate bins that were then consolidated using the MetaWRAP bin refinement module. Subsequently, the refined bins were reassembled using SPAdes v3.10.1 [18]. CheckM v1.0.7 [19] was run on the final reassembled bins to prepare a report of completeness and contamination. Metagenome-assembled genomes (MAGs) with more than 90% completion and less than 5% contamination were selected for further analysis. For the estimation of the abundance of each bin, the Quant bins module from the MetaWRAP pipeline was used with default parameters.

### 2.4 Diversity and differential abundance analysis

After removing reads aligned to human chromosomes, the filtered reads were assigned taxonomy using the Kraken nt database of KRAKEN2 v2.1.3 [20] (downloaded in May 2023). Species-level taxonomy was used for subsequent analysis. The abundance estimation was performed using BRACKEN v2.9 [21]. The abundance table, taxa table, and metadata were integrated into a single object using the *phyloseq* package in R v4.3.3 for further diversity analysis [22].

CCA and CAP ordinations were used to understand the effect of heavy metals, temperature, pH, and the zone from which the sample was isolated on the microbial composition of the water samples. Beta diversity was evaluated using the Principal Coordinate Analysis (PCoA) technique. The analysis was performed using the ordinate function from the *phyloseq* package. Temperatures were categorized into Low (25.61±0.11°C) and High (31.55±0.24°C). Beta dispersion analysis using the *betadisper* function, followed by ANOVA, was used to infer differences in the variability of the microbial community composition between the temperature groups.

To further investigate the influence of temperature on microbial composition, the ANOSIM (Analysis of Similarities) test was implemented in the vegan R package, with 999 permutations to obtain significance values. An R statistic greater than zero indicates greater between-group than within-group dissimilarity, with statistical significance assessed via permutation. The percentage of microbiome variation explained by temperature was quantified using *Adonis2*. The PERMANOVA Analysis (PERmutational Multivariate ANalysis Of VAriance) was performed using the Bray-Curtis distance matrix as input. The significance of group differences was obtained by a permutation test with 999 permutations. Feature selection using Boruta identified taxa at the species level that differed significantly between high and low temperatures [23]. Additionally, ANCOM-BC was employed to detect taxa with differential abundance across the two temperature conditions [24]. P-values were adjusted for multiple comparisons with the Holm method. Taxa with a q-value below 0.05 were considered significantly differentially abundant.

Shannon and observed alpha diversity indices were measured using the *estimate richness* function of *phyloseq*. The Wilcoxon test was used to evaluate the significance of differences between temperature groups, and the Kruskal–Wallis test for the significance of differences between zones.

### 2.5 Taxonomy and functional classification

The generated MAGs were classified using the classic workflow (*classify wf*) from GTDB-Tk v2.3.2 [25]. The reference database used for classification was R214. To generate a phylogenetic tree, GToTree v1.8.4, a command-line tool based on the Hidden Markov Model (HMM) that builds a tree using a set of singlecopy marker genes, was employed [26]. The default cutoff was the presence of at least 50% of the marker genes. However, since some MAGs did not have 50% of the marker genes, the cutoff was lowered to 20%. The output tree from GToTree was passed to IQ-TREE for bootstrapping for 1000 iterations, and the output from IQ-TREE was visualized using iTOL v7 [27, 28]. For functional annotation, Prokka v1.14.6 was used, which utilises Prodigal for gene prediction [29]. The Prokka output was subsequently used to identify Clusters of Orthologous Genes (COGs) using *cogclassifier* v1.0.5 [30].

### 2.6 Detection of antimicrobial resistance genes

The identification of antimicrobial resistance genes was performed using two approaches. First, the filtered reads in each sample were analysed using the RGI (Resistance Gene Identifier) v6.0.3 tool’s BWT (Burrows-Wheeler Transform) feature, with the default kma aligner [31]. The reads were mapped to the CARD (Comprehensive Antibiotic Resistance Database) v3.2.7 [32]. To ensure robust gene selection, the results were filtered to include only genes with at least 100 mapped reads and an average percentage coverage of 80% or greater. If a read was assigned to multiple reference sequences, the median reference sequence was used to ensure a more accurate representation. RPKM (Reads Per Kilobase Million) was calculated for genes that met these criteria for normalisation across samples.

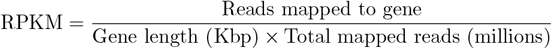

The prediction of ARGs was also implemented by screening the MAGs obtained against the CARD database using the RGI main feature [32]. ORFs with 60% identity with CARD references were further selected for analysis. Furthermore, potential Virulence Factor Genes (VFGs) in MAGs were identified using BLASTp v2.12.0 against the VFDB database (core protein dataset), with a query coverage threshold of 80%, an identity cutoff of 50%, and an e-value cutoff of 1e-5 [33].

### 2.7 Water quality and physicochemical parameters

To investigate water quality, analysis of heavy metals such as magnesium (Mg), manganese (Mn), copper (Cu), arsenic (As), cadmium (Cd), mercury (Hg) and lead (Pb), along with pH and temperature, was carried out using the ICP-MS, Agilent, and Hanna 9829 multiparameter meter, respectively. The findings were compared with the guidelines prescribed by the WHO (2022) and IS 10500:2012 drinking water quality standards (**Table 1**).

**Table 1:** Heavy metal analysis and their comparison with permissible limits prescribed by the WHO, 2022 and IS:10500:2012.

| S.no. | Samples | Metals/Parameters | Observed value(ppb) | WHO Limit (2022)<br>(ppb) | IS:10500:2012<br>Limit (ppb) |
| --- | --- | --- | --- | --- | --- |
| 1. | W1-W21 | Arsenic (As) | 0.56-3.05 | 10 | NP<sup>1</sup> |
| 2. | W1-W21 | Cadmium (Cd) | 0-0.11 | 3 | 3 |
| 3. | W1-W21 | Copper (Cu) | 0.25-11.60 | 2000 | 50 |
| 4. | W1-W21 | Lead (Pb) | 0-1.72 | 10 | 10 |
| 5. | W1-W21 | Manganese (Mn) | 0.56-105.19 | 80 | 100 |
| 6. | W1-W21 | Mercury (Hg) | 0-0.14 | 6 | 1 |
| 7. | W1-W21 | Magnesium (Mg) | 4391.08-28792.60 | NP<sup>1</sup> | 30000 |
| 8. | W1-W21 | Zinc (Zn) | 0-306.99 | NP<sup>1</sup> | 5000 |
| 9. | W1-W21 | Iron (Fe) | 0-277.42 | NP<sup>1</sup> | 300 |
| 10. | W1-W21 | pH | 6.80-7.50 | NP<sup>1</sup> | 6.5-8.5 |
| 11. | W1-W21 | Temperature °C | 25.4-32.0 | NP<sup>1</sup> | NP<sup>1</sup> |

<sup>1</sup>NP: Not provided

## 3 Results

### 3.1 Water quality (heavy metals, pH, and temperature)

The pH and temperature of the drinking water samples (W1–W21) ranged from 6.8 to 7.5 and 25.4°C to 32°C, respectively. The pH levels were within the acceptable limits of IS:10500:2012 for potable water. Furthermore, toxic heavy metals with known carcinogenic potential: arsenic (As), cadmium (Cd), lead (Pb), copper (Cu), and mercury (Hg) were detected at concentrations of 0.56–3.05, 0–0.11, 0–1.72, 0.25–11.6, and 0–0.11 µg/ml (ppb), respectively. These values comply with the drinking water quality standards set by WHO and IS:10500:2012 (**Table 1**). Essential minerals such as magnesium (Mg), manganese (Mn), iron (Fe), and zinc (Zn) were found to be within the permissible limits outlined by these standards. Thus, it can be inferred that in terms of physicochemical contaminants, the drinking water obtained from the metropolitan city falls under the criteria “Safe for drinking purposes”.

### 3.2 Diversity of drinking water metagenome

The taxonomy assignment of the filtered reads revealed that, on average, 59.19 ± 6.34% of the reads in a sample were successfully classified, with 40.85 ± 6.35% classified at the species level (**Supplementary Table 1**). Only reads assigned to Bacteria, Archaea, Fungi, and Viruses were included in the diversity analyses. Bacteria dominated the microbial composition, comprising 98.41% of the identified taxa, while the other domains accounted for 0.38%, 0.99%, and 0.21%, respectively. Further, to identify the core taxa present across all samples, species prevalence was calculated at a read cutoff of more than 10,000, revealing five species with 100% prevalence: *Afipia carboxidovorans, Klebsiella pneumoniae, Pseudomonas aeruginosa, Sphingopyxis macrogoltabida*, and *Variovorax paradoxus*. When the read cutoff was reduced to 1,000, the number of species with a prevalence of 100% increased to more than 500. The composition of Bacteria and Archaea at the Class level, followed by a plot of the abundances of the 15 most abundant bacterial/archaeal species, is shown in (**Fig. 1**). At the species level, bacteria such as *Afipia carboxidovorans, Bosea sp*., *Bradyrhizobium sp*., *Hydrogenophaga crocea, Limnobacter sp*., *Mycolibacterium fluoranthenivorans, Mycolibacterium gilvum*, and *Pseudomonas alcaligenes* dominated the community structure. In contrast, the fungal community was summarised at the order level, and the 15 most abundant species were visualised in a separate plot. The community was characterised by the prevalence of species such as *Amniculicola longissima, Epicoccum nigrum, Fusarium oxysporum, Lasiodiplodia theobromae, Leptosphaeria biglobosa, Moeziomyces antarcticus*, and *Neofusicoccum parvum*. The composition at the order level, along with the abundances of the top 15 species, is shown in **Fig. 2**.

**Fig. 1:**
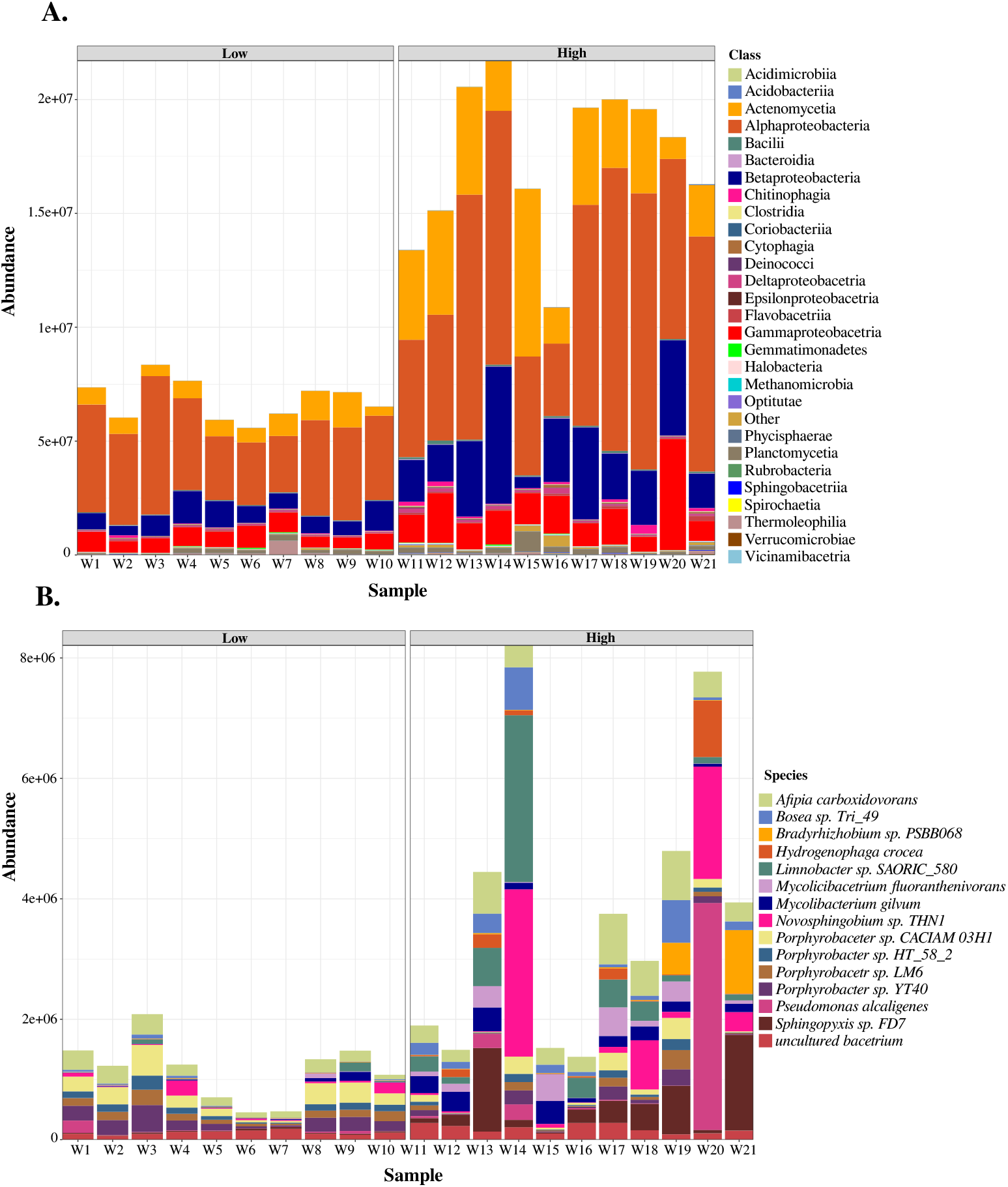
**A**.Bacteria/Archaea composition at class level, detected above 1000 reads in all samples. **B**. Distribution of the 15 most abundant bacterial/archaeal species across samples.

**Fig. 2:**
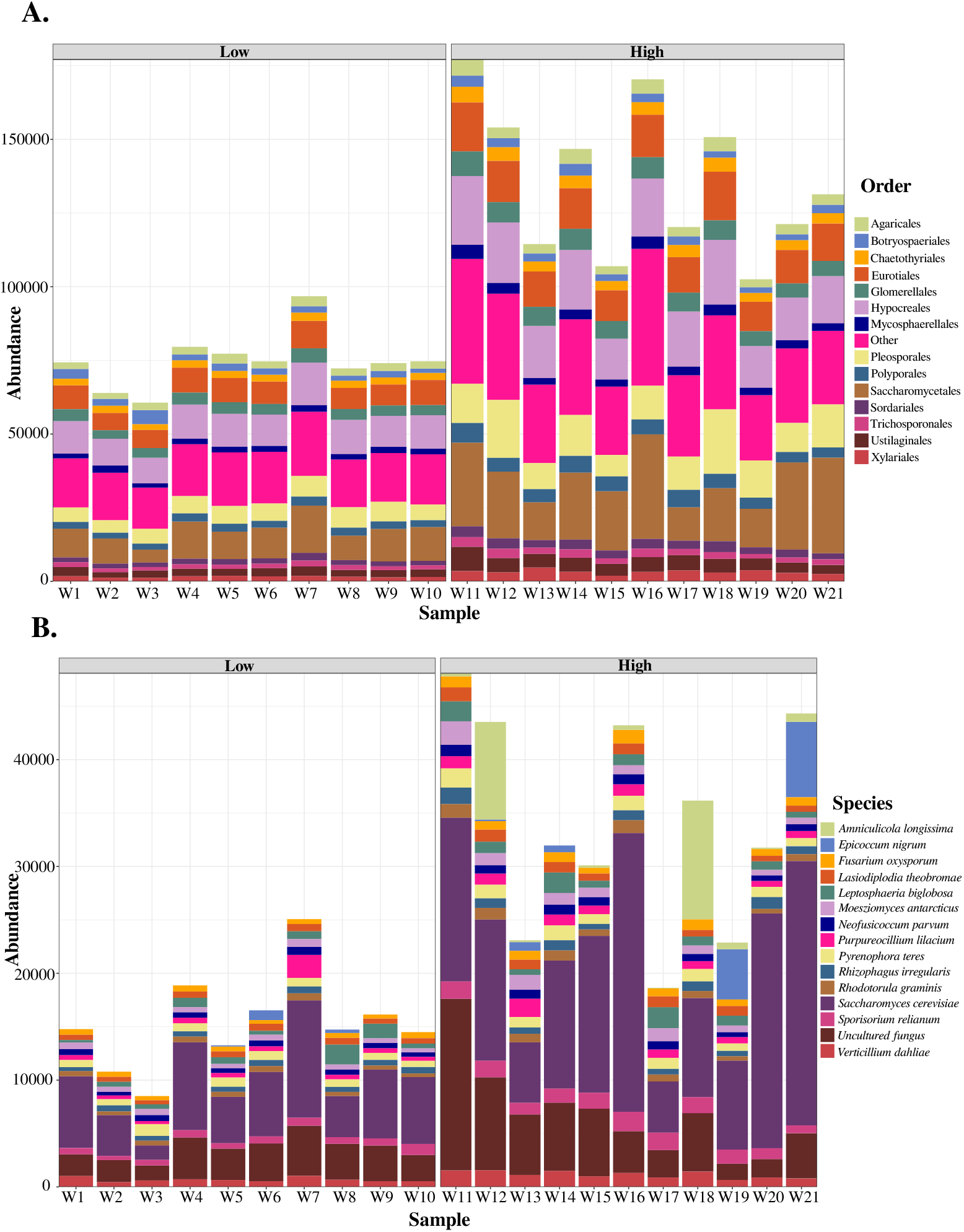
**A**. Fungal community composition at the order level, detected above 1000 reads in all samples. **B**. Distribution of the 15 most abundant fungal species across samples.

### 3.3 Drivers of microbial diversity

The effects of covariates on the composition are shown in Figure (**Fig. 3**). We observe that the temperature and the mercury content are correlated. Since mercury levels have a small range and are well within permissible limits, we infer that temperature appears to drive the microbial composition. Although, pH also appears to influence the microbial composition, its range is limited, and the results are inconsistent, with PERMANOVA findings contradicting those of CCA. Similar inconsistencies are observed for heavy metals.

**Fig. 3:**
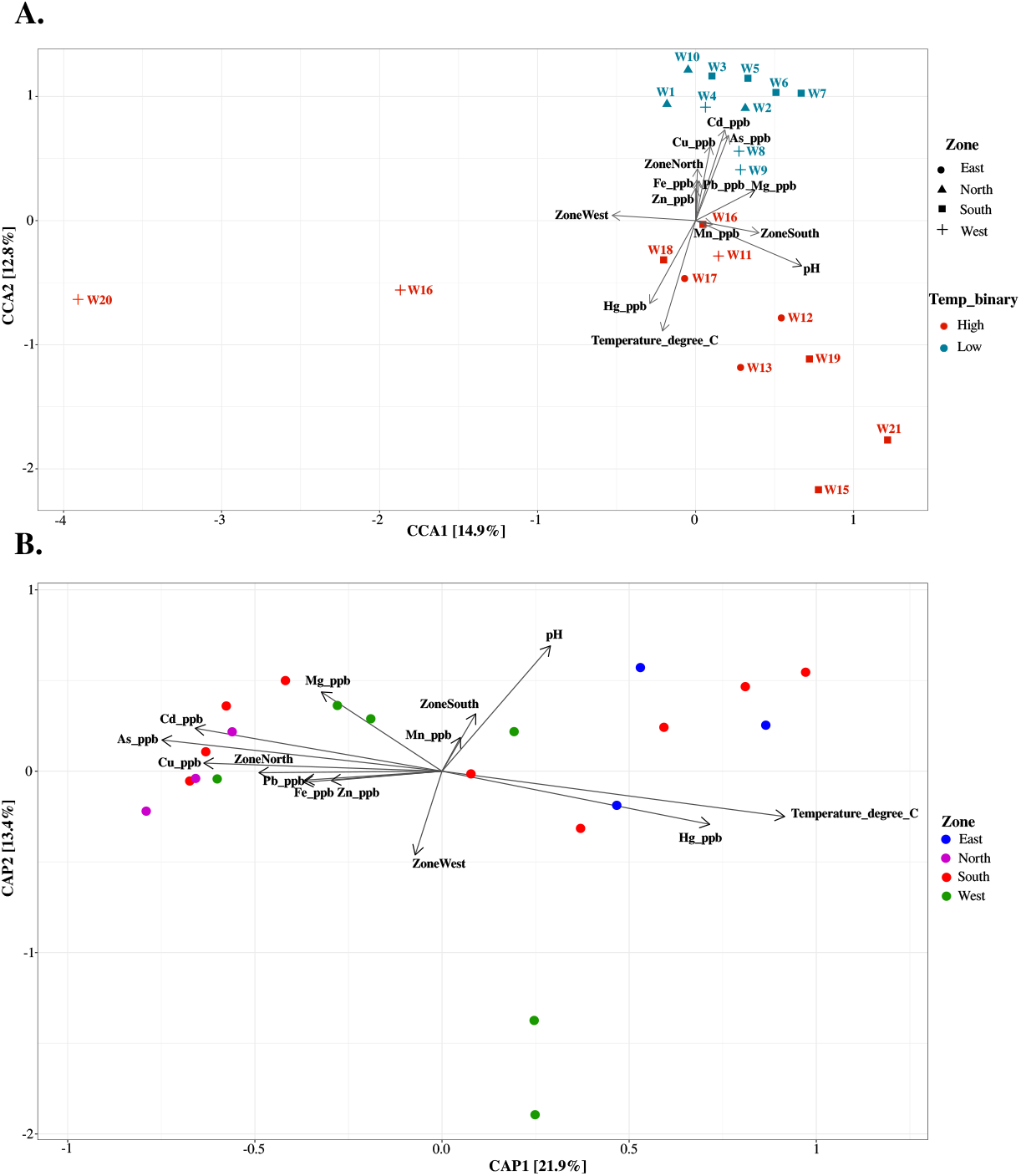
Canonical Correspondence Analysis (CCA) and Canonical Analysis of Principal Coordinates (CAP) plots showing the relationship between microbial communities and environmental variables.

Across all analyses, changes in temperature consistently correlate with shifts in the microbial composition. Therefore, we focus on analysing the effects of temperature on microbial diversity in more detail.

Principal Coordinates Analysis (PCoA) shows the clustering of samples from the low and high temperature groups (**Fig.4B**). The sample sizes were balanced, with 10 samples from the lower temperature group (25°C) and 11 from the higher temperature group (32°C). The homogeneity of the variances was assessed using Beta Dispersion with the *betadisper* function. ANOVA revealed significant differences in beta dispersion between groups (p = 0.002413), indicating variability in the composition of the microbial community under different temperature conditions **(Supplementary Fig. S1)**.

The effect of temperature on the diversity of the microbial community in the samples was studied **Fig 4**. PERMANOVA, performed using the Adonis2 package, also identified temperature as a significant contributor to microbial diversity, with an R^2^ value of 0.4265 and p = 0.001. However, since PERMANOVA results can be influenced by differences in beta dispersion, we performed an ANOSIM analysis. ANOSIM confirmed greater similarity within the temperature groups than between them (p = 0.001, R = 0.4513), further supporting the role of temperature in shaping the composition of the microbial community.

**Fig. 4:**
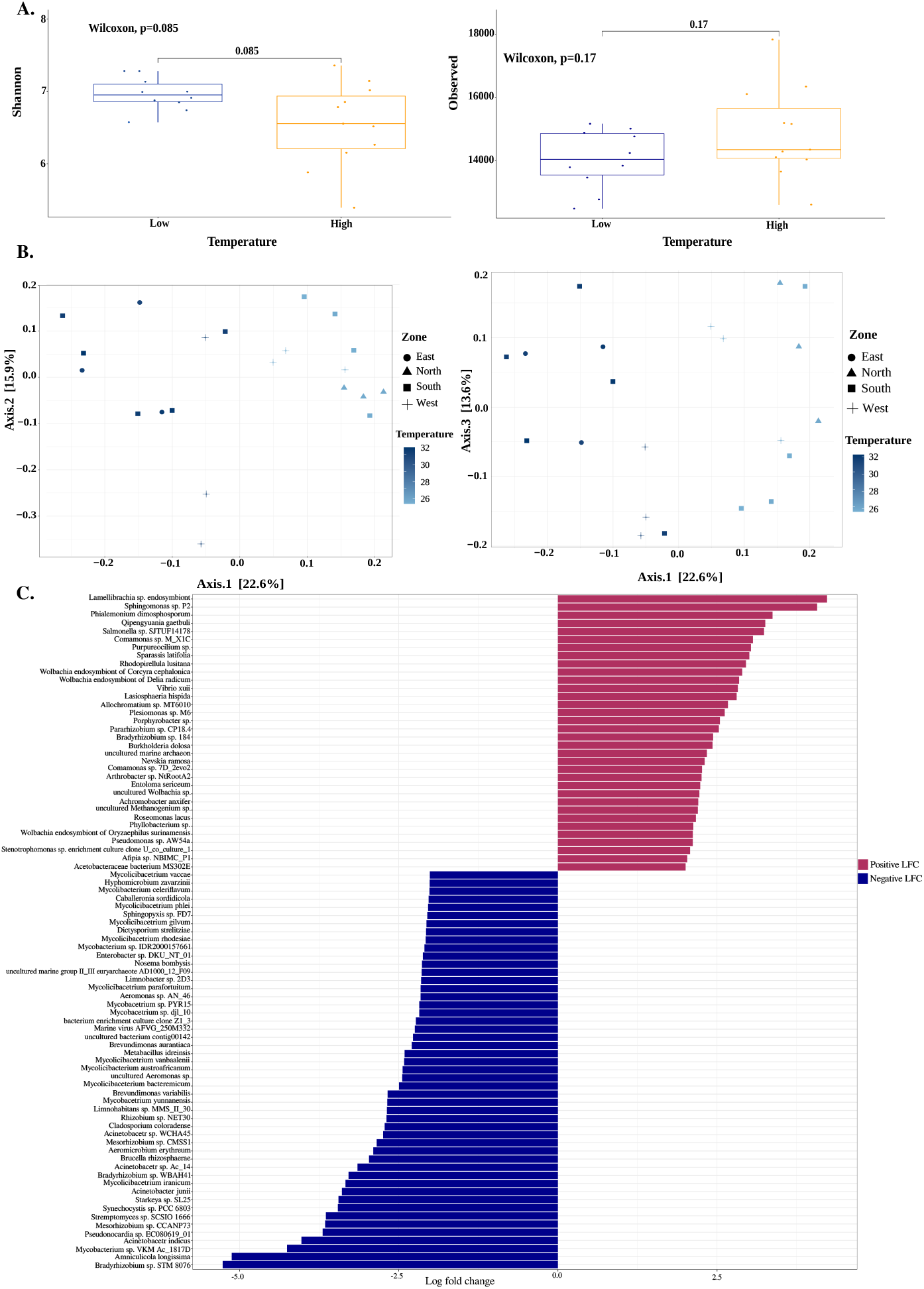
Effect of temperature on the diversity of the microbial community in the samples **A**. Alpha diversity was estimated between high and low-temperature groups using the Shannon and Observed diversity indices. The p-values for the Shannon index (0.085) and the Observed species index (0.17) indicate that the differences in species richness and evenness between the two temperature groups were not statistically significant. **B**. Beta diversity comparisons were visualized using Principal Coordinates Analysis (PCoA). These plots reveal that, while species richness and evenness do not differ significantly between temperature groups, temperature plays a crucial role in determining the species composition of the samples. **C**. Differential abundance analysis of taxa at high and low temperatures. Red represents taxa more abundant in low temperatures

Alpha diversity was measured using both the Shannon index (which measures richness and evenness) and the observed species index (which estimates richness) to estimate the microbial richness and evenness across zones **Fig. 4A**. No significant differences were observed, which aligns with expectations, as the water supplied to the zones was subjected to uniform treatment. Although the PCoA plot shows a separation between the North and East zones, their species richness and evenness were not significantly different. In contrast, the North and West zones were clearly separated in the PCoA plot, with a significant difference in alpha diversity (p = 0.024) (**Supplementary Fig. S2**).

To identify differentially abundant taxa under different temperature conditions, we used a random forest-based wrapper, Boruta, to the filtered *phyloseq* object (prevalence = 0.1, detection = 50). We also used ANCOM-BC on the filtered data, and the results were compiled with a significance cutoff of 0.05 with the p adjustment method - “holm.” The taxa identified by ANCOM-BC are presented in **Fig. 4C**. In particular, both ANCOM-BC and Boruta identified common taxa that were more abundant at high temperatures, including *Acinetobacter indicus, Acinetobacter junii, Acinetobacter sp*. WCHA45, *uncultured bacterium* contig00142, *Hyphomicrobium zavarzinii*, and *Acinetobacter sp*. SCLZS86. A detailed list of taxa identified by BORUTA and ANCOM-BC is presented in **Supplementary Table 2**

### 3.4 Community structure of drinking water

After applying a stringent filter (over 90% completion and less than 5% contamination), 358 bacterial MAGs were obtained for further analysis, with 233 bins from high temperature and 125 from low temperature. dRep was not run to preserve information regarding sample-specific genomic content of the individual strains for COG analysis **(Fig. 5)**. Classification of the MAGs revealed that Pseudomonadota was the most dominant phylum in 68.44 ± 14.35% of the samples in various zones. This was followed by Planctomycetota at 10.74 ± 8.68%, Bacteroidota at 6.35 ± 6.28%, and Actinomycetota at 6.92 ± 6.80%. Pseudomonadota was observed to be the dominant phylum in all zones, and Actinomycetota was not observed in the north zone. For a deeper understanding of the microbial composition, the MAGs were quantified to find the most abundant groups. *Sphingomonadaceae* was the most abundant family in the community. The abundance of different families is shown in **Supplementary Fig. S3**.

**Fig. 5:**
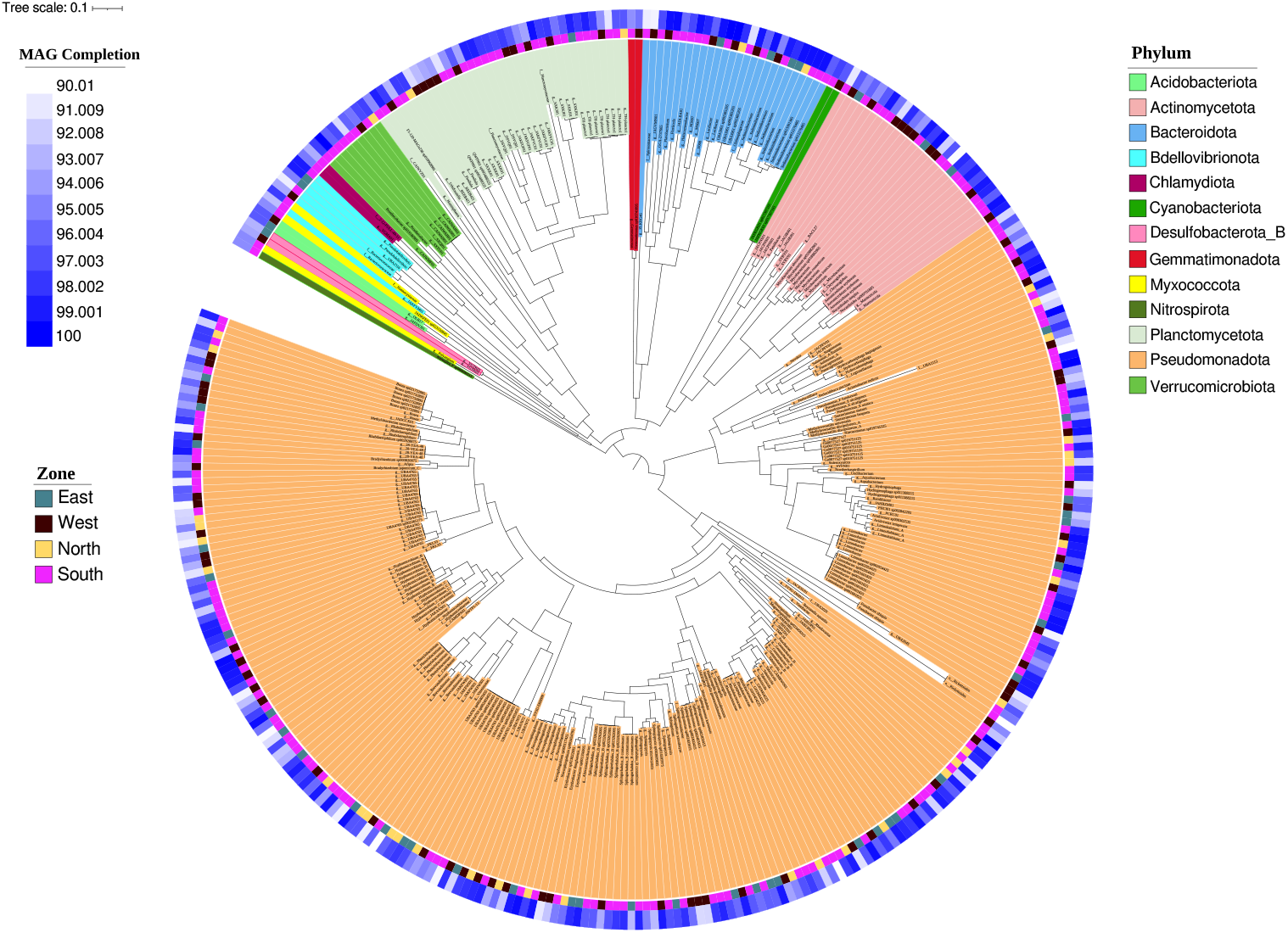
Phylogenetic tree of the MAGs. Phylum-level classification of the MAGs obtained from the samples, showing their completion levels and the respective zones from which the samples were collected.

We used GRiD to identify genomes with higher replication rates in the samples, as a substitute for viability analysis. GRiD scores identify actively dividing bacterial cells by computing the ratio of the origin of replication to termination sites. Among the families analysed, *Burkholderiaceae B* exhibited the highest growth rate. Interestingly, species belonging to the *Sphingomonadaceae* and the *Burkholderiaceae B* families demonstrated high growth rates and substantial abundance within the community (**Fig. 6, Supplementary Fig. S4**).

**Fig. 6:**
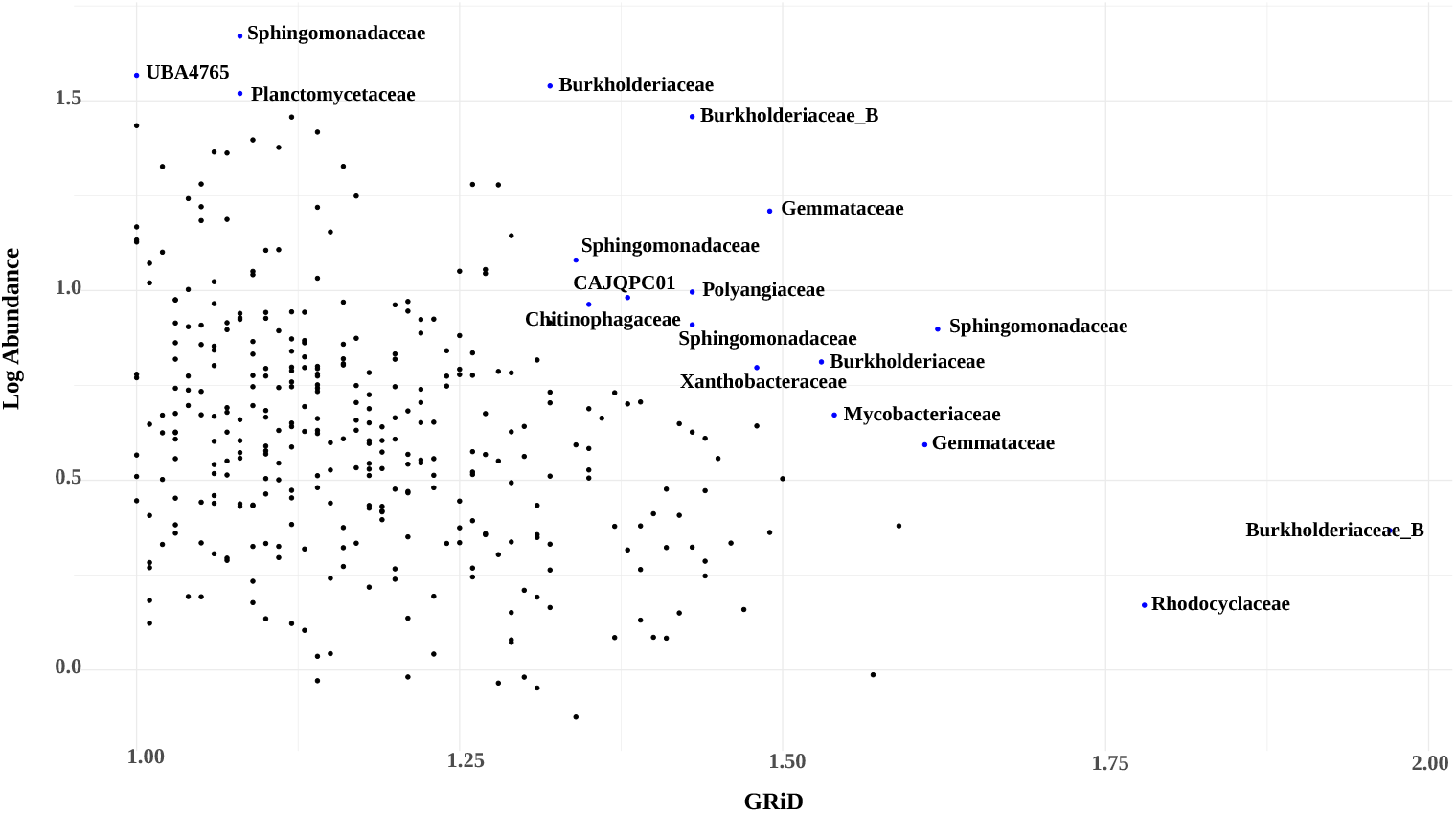
Replication rate of families identified in the samples. Scatter plot depicting bacterial families with a GRiD score greater than 1.3 (indicating actively dividing bacteria) and a log-transformed abundance exceeding 0.75. Only families meeting these thresholds are labeled.

Our primary objective was to gain a broader understanding of the microbes present in drinking water. We annotated Open Reading Frames (ORFs) to identify coding sequences (CDS). A total of 969,369 distinct Clusters of Orthologous Genes (COGs) were identified, belonging to 25 functional categories. To ensure specificity and sensitivity, we applied a 30% identity cutoff, which refined the number of COGs to 661,322. A comparative analysis of these COGs in all zones revealed that the number of genes associated with different functional categories did not differ significantly. The abundance of COG categories is represented in **Supplementary Fig. S5**.

Although the concentrations of all metals in the samples were within the permissible limits, the concentration of iron in one of the samples was quite different from the mean values. Our analysis identified three COGs that are associated with iron metabolism: COG0735 (Fe^2+^ or Zn^2+^ uptake regulation protein Fur/Zur), COG1914 (Mn^2+^ or Fe^2+^ transporter, NRAMP family), and COG0672 (High-affinity Fe^2+^/Pb^2+^ permease). Interestingly, the presence of COG0735 was also high in the *Sphingomonadaceae* family and samples at high temperatures (**Fig. 7, Supplementary Fig. S5**). A complete presence/absence list of all COGs identified in the MAGs has been presented in **Supplementary Table 2**.

**Fig. 7:**
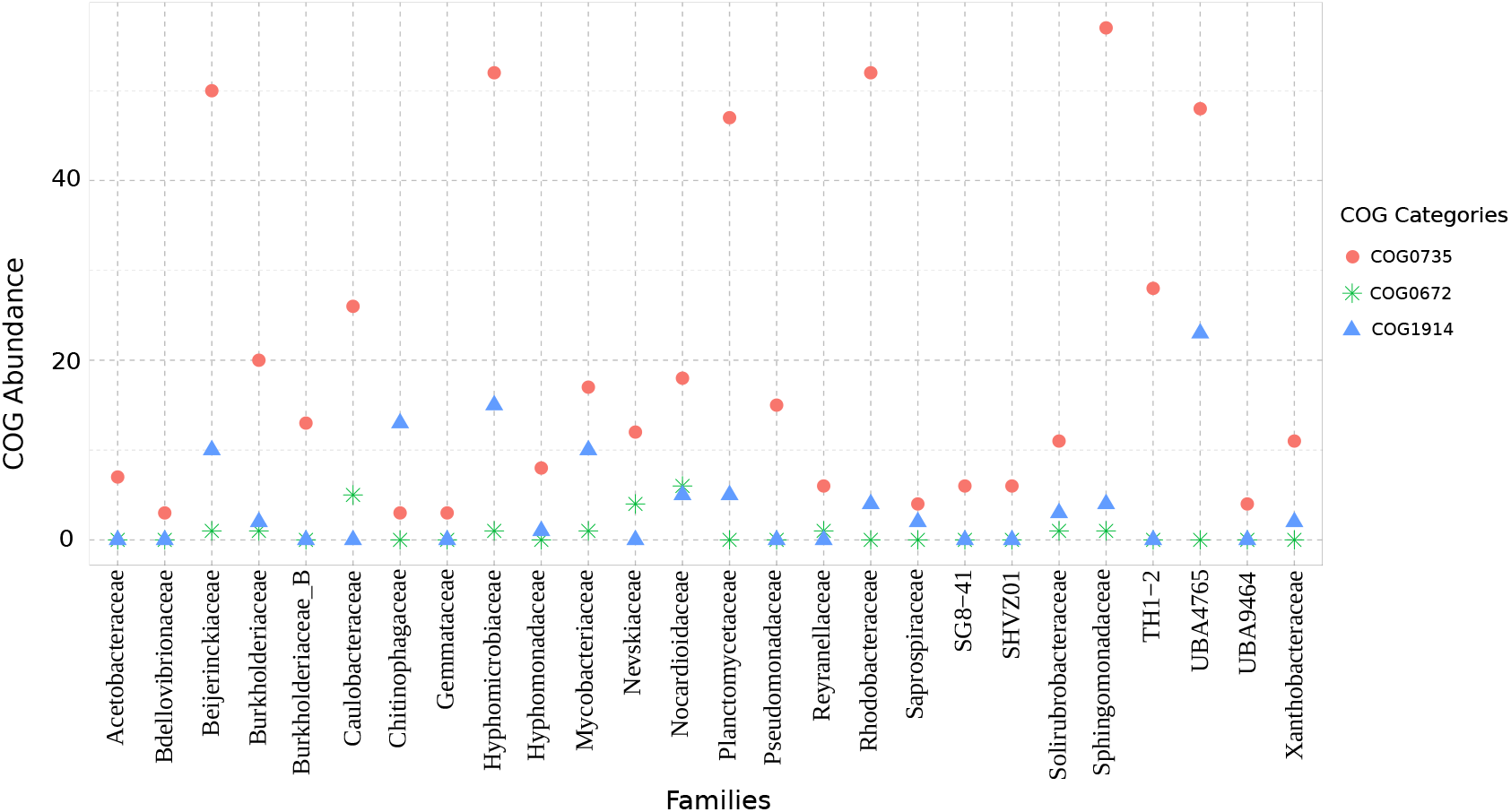
Distribution of COGs involved in iron metabolism among the bacterial families. COG0735 (Fe^2+^ or Zn^2+^ uptake regulation protein Fur/Zur), COG1914 (Mn^2+^ or Fe^2+^ transporter, NRAMP family), and COG0672 (High-affinity Fe^2+^/Pb^2+^ permease). Only families with more than 10% relative abundance are shown in the figure.

### 3.5 Distribution of ARGs in the drinking water

ARGs were characterised from metagenomic reads using RGI-bwt, following the filtration criteria described in the Methods section. We identified 82 unique ARGs that span 43 different families of AMR genes after applying these filters. Normalisation was carried out by calculating the RPKM for each gene identified to quantify the abundance of antibiotic resistance genes and comparing them between different samples. In particular, genes associated with resistance to aminoglycoside, elfamycin, fluoroquinolone, macrolide, and tetracycline drug classes were consistently detected in all samples and zones (**Fig. 8**). Among all drug classes, aminoglycoside and macrolide antibiotics had the highest diversity of ARGs, with 22 and 25 unique genes, respectively.

**Fig. 8:**
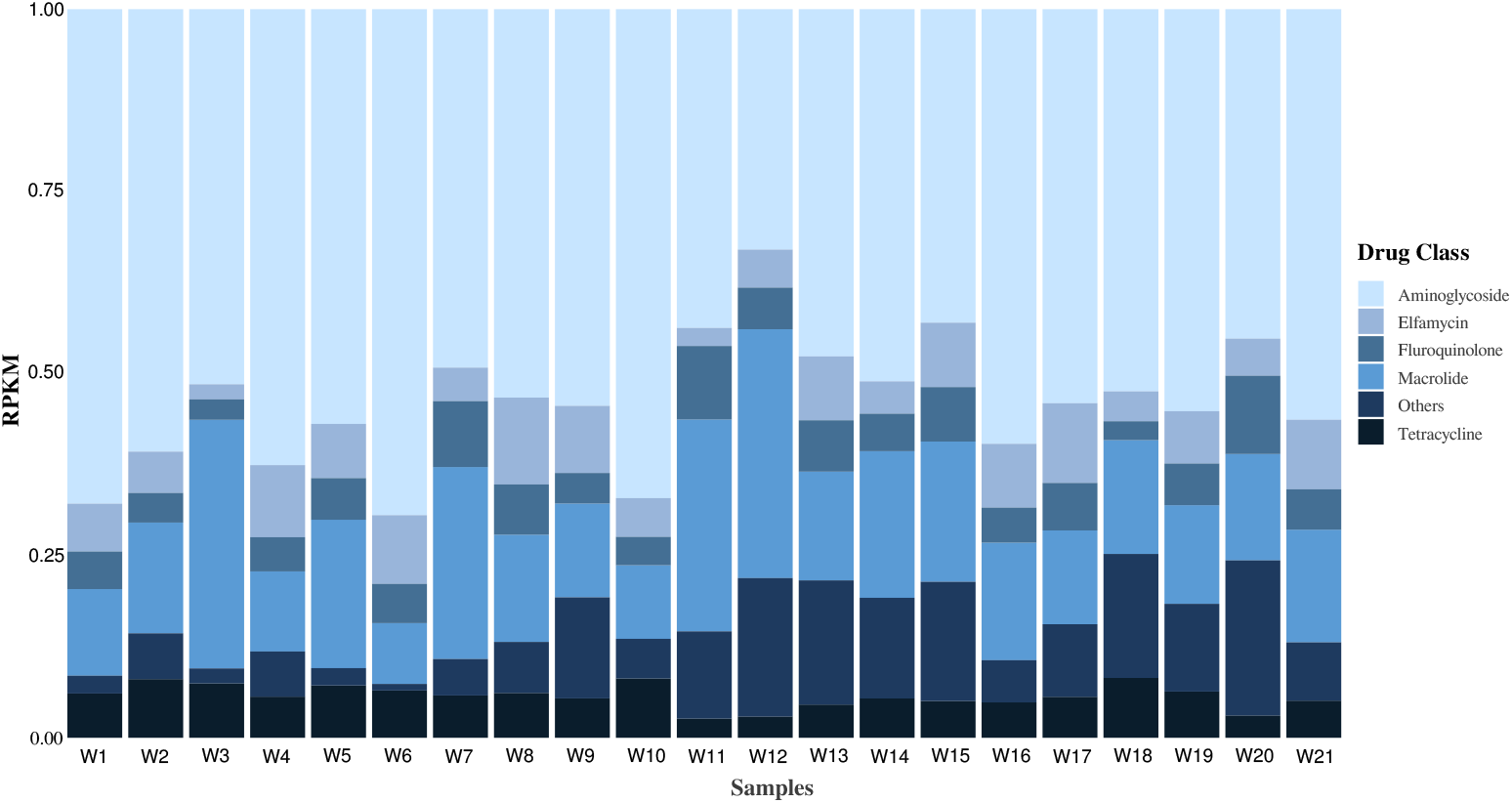
Stacked bar plot showing the abundance of antimicrobial resistance genes (ARGs) conferring resistance to different drug classes across samples. The x-axis represents individual samples, while the y-axis indicates the normalized abundance of ARGs expressed in RPKM (Reads Per Kilobase of transcript per Million mapped reads). Each stack corresponds to a specific drug class, with the height proportional to the total abundance of ARGs identified for that class. Only the top five drug classes for which ARGs are identified in all samples are shown here.

Multidrug-resistant genes comprised 14% of all the genes identified. The *Pseudomonas aeruginosa* CpxR gene, detected exclusively in the W18 sample, showed resistance to 15 drug classes, the highest among all genes identified. In general, the W18 sample contained resistance genes for 28 of the 29 drug classes identified. Resistance to the drug classes aminocoumarin, monobactam, and sulfonamide was also detected exclusively in this sample. **Supplementary Table 2** provides a detailed list of drug classes, their associated resistance genes, and their corresponding RPKM values.

The mechanisms by which genes in this study confer resistance include antibiotic efflux, antibiotic inactivation, target replacement, target alteration, and resistance by absence. Although most genes conferred resistance through target alteration, most multidrug resistance genes conferred resistance primarily through the efflux mechanism.

To better understand ARGs in metagenome-assembled genomes (MAGs), we used the rgi-main tool with the CARD database. The adeF gene, which confers resistance to fluoroquinolone and tetracycline, was the most prevalent ARG in the samples, detected in 15 bacterial families, with the highest representation in the *Pseudomonadaceae* family. which also uniquely harbored the multidrug resistance gene YajC. It exhibited resistance to ten drug classes, including fluoroquinolones, cephalosporins, glycylcyclines, penams, tetracyclines, oxazolidinones, glycopeptides, rifamycins, phenols, disinfectant agents, and antiseptics. Furthermore, the identified Pseudomonas aeruginosa soxR gene was resistant to 8 drug classes and was found exclusively in *Xanthobacteraceae* (**Fig. 9, 10**). Among drug classes, fluoroquinolones had the highest prevalence of resistance-associated genes in MAGs. A detailed list of genes identified in MAGs, along with their mechanism of action and drug classes to which they confer resistance, is presented in **Supplementary Table 2**. In general, most ARGs were found in the *Pseudomonadaceae* and *Mycobacteriaceae* families. The distribution of ARGs across bacterial families is shown in **Fig. 10**.

**Fig. 9:**
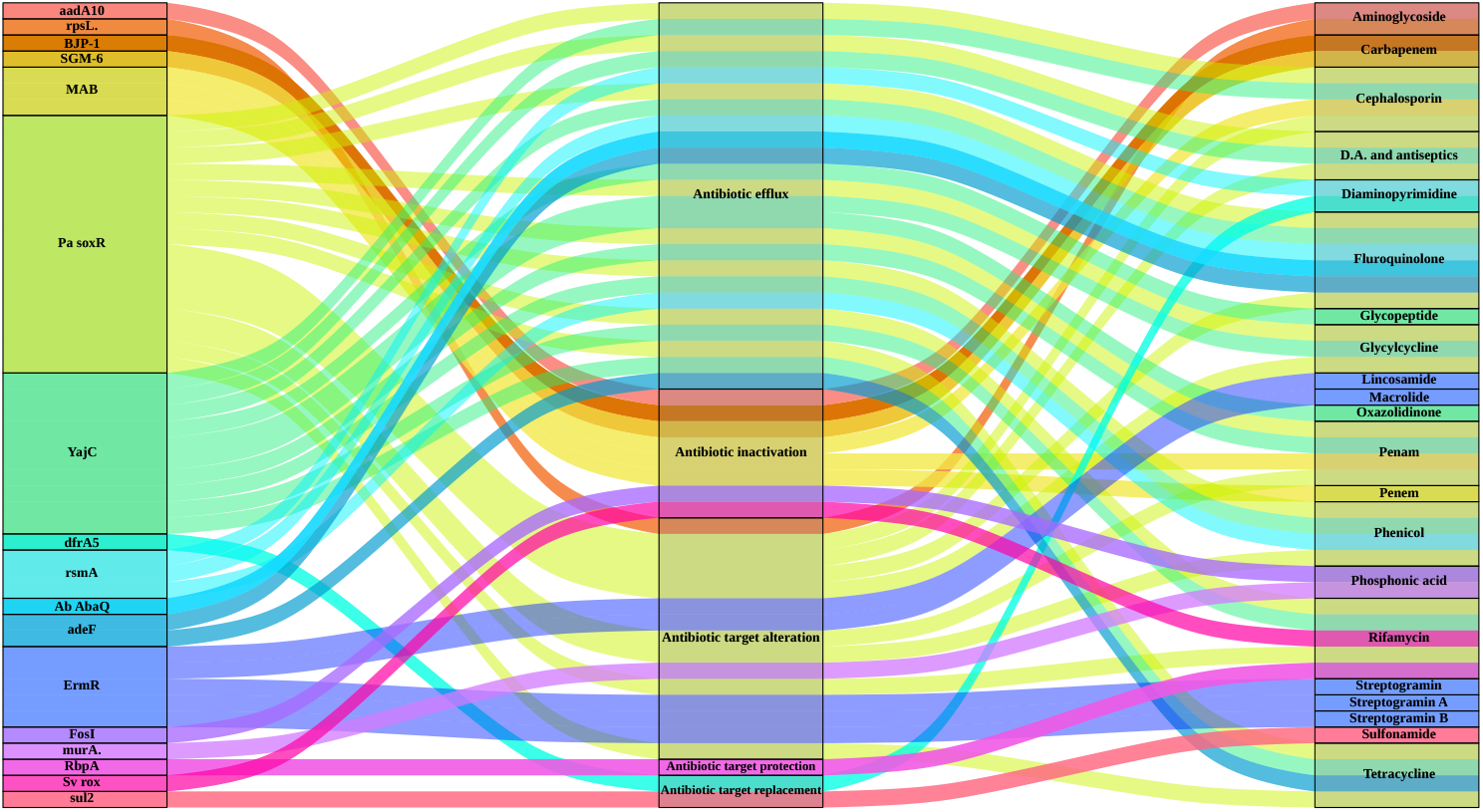
Genes detected in the MAGs, their mechanism of action, and the drug classes to which they are resistant. Amongst all ARGs represented here, Pa soxR, MAB, YajC, rsmA, adeF, and ErmR are multidrug resistant. The abbreviations used in the figure are as follows: Pa soxR: *Pseudomonas aeruginosa* soxR, Ab abaQ: *Acinetobacter baumannii* abaQ, murA: *Mycobacterium tuberculosis* intrinsic murA conferring resistance to fosfomycin, rpsL: *Mycobacterium tuberculosis* rpsL mutations conferring resistance to streptomycin, D.A.: Disinfecting agents, Sv: *Streptomyces venezuela* rox.

**Fig. 10:**
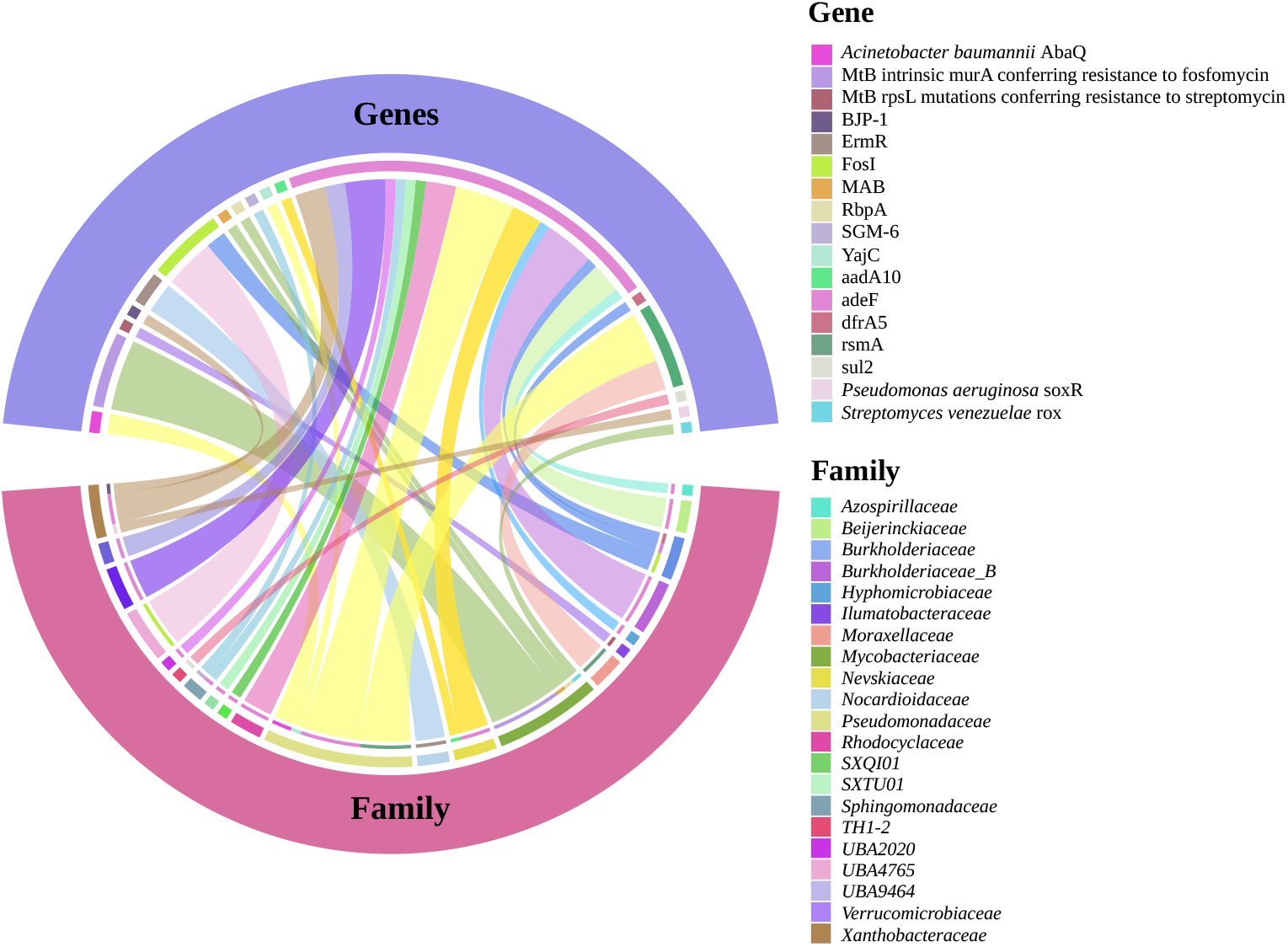
Distribution of genes identified in the MAGs across different Families. The gene sectors are displayed from left to right in the same order as they appear in the legend. Similarly, the Family sectors are displayed from right to left, maintaining the order specified in the legend.

### 3.6 Virulence Factors identified in Drinking water

The virulence factors obtained after filtration belonged to 14 categories whose distribution is shown in **Supplementary Fig. S6**. Our analysis indicates that nutritional/metabolic factors are prevalent in all families, closely followed by immune modulation and motility genes. Given that the *Sphingomonadaceae* family was the most abundant, as determined from our MAG classification, it also exhibited the highest number of virulence factors. Consequently, no definitive conclusions can be drawn about the enrichment of specific genes in one family over another.

## 4 Discussion

Periodic monitoring of drinking water supplied through the public water distribution system post-treatment is essential. This ensures that quality is maintained and that public health is not adversely impacted by persistent abiotic or biotic contaminants. The emergence of metagenomics offers a comprehensive and holistic understanding of the microbial community that will help develop effective management and mitigation strategies to ensure safe and high-quality drinking water [34].

In the present study, we sampled water from 21 locations in four zones of an Indian city. To our knowledge, this is the first metagenomic study of drinking water from India. The most abundant phyla in our analysis were Pseudomonadota (Proteobacteria), Planctomycetota, Bacteroidota, and Actinomycetota. This finding is consistent with previous reports [35].

Diversity analyses showed that temperature was the main factor that influenced the microbial composition, followed by zone to a lesser extent. The impact of metal concentration on microbial diversity could not be assessed due to the narrow range of values for some metals and the limited sample size, which makes it difficult to draw any conclusions. Additionally, there was at least one outlier for each metal that would have affected the analysis, so we focused our study on temperature and zone. The levels were also within acceptable limits, making further analysis unnecessary.

While PMA treatment is often recommended for viability assessment, we chose not to apply it in this study. This decision was based on recent evidence showing that PMA, though effective in suppressing DNA from dead cells, also leads to a significant reduction in total DNA yield across all taxa, including viable ones. This reduction can limit sequencing depth and sensitivity, particularly in oligotrophic environments such as potable water. Given our aim to broadly profile microbial communities across diverse zones and temperature gradients, we prioritized maintaining DNA yield to capture a more comprehensive taxonomic landscape [36].

Among the microbial families identified in our samples, *Sphingomonadaceae* was the most dominant, with 45 MAGs assigned to this family. Members of this family have been identified in various ecosystems, including soil, corals, clinical samples, and drinking water [37]. They are also known to be strictly aerobic chemoautotrophs and play a crucial role in initiating biofilm formation and contribute significantly to biofilm maturation [38, 39]. Furthermore, the higher prevalence of the Fur gene within this family highlights intriguing avenues for future research. The Fur gene regulates cellular iron homeostasis, with the iron-Fur complex activating in the presence of elevated ferrous iron levels, thus repressing iron metabolism [40]. However, no correlation was found between the abundance of the *Sphingomonadaceae* family and the concentration of iron in our study. While a trend indicated higher levels of COG0735 in the high-temperature group, the relationship remains inconclusive due to the limited sample size.

In addition to members of the *Sphingomonadaceae* family, members of the *Rhodobacteraceae* and *Beijerinckiaceae* families, although they had half the abundance of *Sphingomonadaceae*, showed a significant presence of COG0735 (**Fig. 7**). A higher abundance of nutritional/metabolic virulence factors in *Rhodobacteraceae* and *Beijerinckiaceae* highlights their potential roles in competitive nutrient acquisition and survival under trace metal-limited conditions (**Supplementary Fig. S6**). *Rhodobacteraceae*, in particular, are known for their dependence on trace metals, which supports their diverse metabolic pathways and ecological strategies [41]. The profiles of virulence factors and COGs in these two families raise important questions about the role of iron in their potential synergistic interactions or competition, which could be explored in future studies.

The analysis of antibiotic resistance genes revealed the presence of the ermR gene in both metagenomic reads and MAGs. The sole carrier of this gene is *Aeromicrobium erythreum*, which was also binned from our samples. The gene was found in samples W11, W18, and W19 (**Supplementary Table 2**). This gene, previously known as ermA, has been reported to confer resistance to macrolide-lincosamide-streptogramin (MLS) antibiotics, consistent with our findings. MAGs for *Aeromicrobium erythreum* were also binned from these three samples: W18 bin.26.orig, W19 bin.11.orig, and W11 bin.8.orig (**Supplementary Table 1**). This bacterium has not been previously reported in drinking water [42, 43]. Macrolides were widely used to treat upper respiratory tract infections (RTIs) generally caused due to bacterial species such as *Streptococcus pneumoniae, Haemophilus influenzae*, and *Streptococcus pyogenes*. Over time, these bacterial species have developed antibiotic resistance through genes such as ermR and ermA, which markedly reduce the therapeutic efficacy of macrolides and consequently necessitate the use of broader-spectrum or more toxic antimicrobials, including fluoroquinolones and tetracyclines [44]. Furthermore, the high prevalence of the adeF gene, as noted earlier (Section 3.5), confers resistance to a wide range of antibiotics, including fluoroquinolones and tetracyclines. The co-occurrence of adeF with ermF may further constrain therapeutic options and facilitate the persistence of high-risk pathogens within the *Pseudomonadaceae* family, thereby intensifying both clinical management challenges and environmental health risks.

While the presence of a gene does not confirm its expression, the detection of ARGs in potable water is a significant health concern. The global spread of ARGs threatens to render common bacterial infections untreatable, and water systems can act as crucial reservoirs and conduits for their dissemination into the human population. This study underscores the critical need for systematic surveillance of ARGs in the drinking water. Future studies should not only identify the presence of these genes but also investigate their expression levels and potential for horizontal gene transfer to pathogenic bacteria. Such research is vital to fully assess the health risks posed by the ARGs to inform public health policy [45].

## 5 Conclusion

A comprehensive understanding of the microbial community of drinking water is crucial to ensure its quality and safety. In this study, we used metagenomics to analyse the microbial diversity of drinking water supplied through the public distribution systems of an Indian city. Our results showed that temperature was the most significant physicochemical factor influencing microbial community composition. The influence of other factors could not be thoroughly assessed due to limited sample size and insufficient variation across samples. While propidium monoazide (PMA) treatment was not applied, and therefore we could not distinguish between viable and nonviable microbes, this choice was made to preserve DNA yield and maximize detection sensitivity, particularly important in low-biomass systems such as potable water. Our analysis also identified antimicrobial resistance genes (ARGs); Although the presence of ARGs does not directly indicate a risk, it underscores the importance of future research focused on horizontal gene transfer and risk assessment of these ARGs. This study reinforces previous observations of drinking water microbiomes while also providing new insights into their composition. It also improves our understanding of the drinking water ecosystem and reiterates the importance of monitoring to protect public health.

## Supplementary information

The following are the Supplementary data for this article.

Supplementary Figures

Supplementary Table 1

Supplementary Table 2

## Acknowledgements

V.K., K.T., and I.T. are thankful to the Director of Zoological Survey of India and the RAMC committee for funding the project titled “Drinking Water Microbiome: Monitoring of the Water Quality and Associated Health Implications.” S.S. acknowledges the half-time research assistantship from the Ministry of Education, Government of India. B.L. acknowledges support from the Centre for Integrative Biology and Systems mEdicine (IBSE). A.R. acknowledges support from the project RB22231279BTHINT008481.

## Declaration of generative AI and AI-assisted technologies in the writing process

During the preparation of this work, the author(s) used the Large Language Model (LLM) tool ChatGPT and Grammarly to assist with grammatical corrections and to improve the readability of the text. After using these tools, the author(s) reviewed and edited the content as needed, and take full responsibility for the final content of the publication.

## Data availability statement

The generated sequences were submitted to NCBI under the bio project ID: PRJNA1008921 with accession number SRR25754722 to SRR25754742.

