## Supplementary File for "Metagenomic Profiling of Municipal Drinking Water Microbiomes in an Indian City: Insights into Diversity, Water Quality, and AMR Potential"

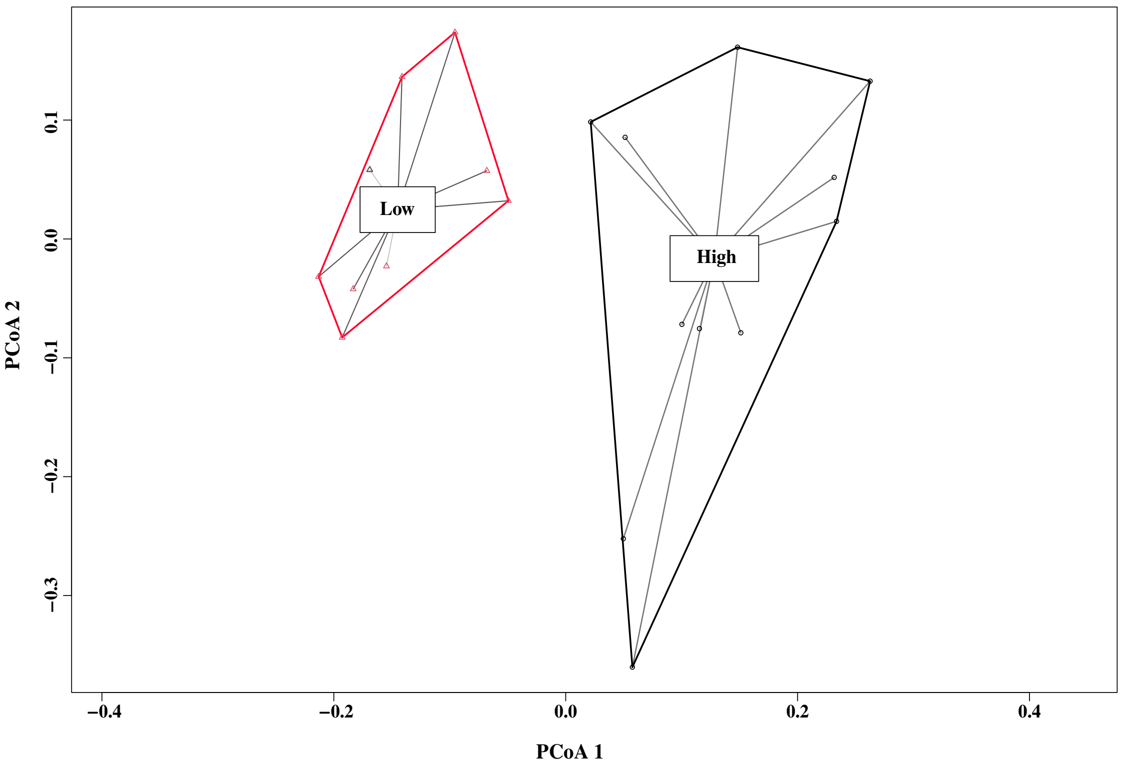


**Fig. S1** Beta dispersion comparing microbial community variation between high and low temperature groups.

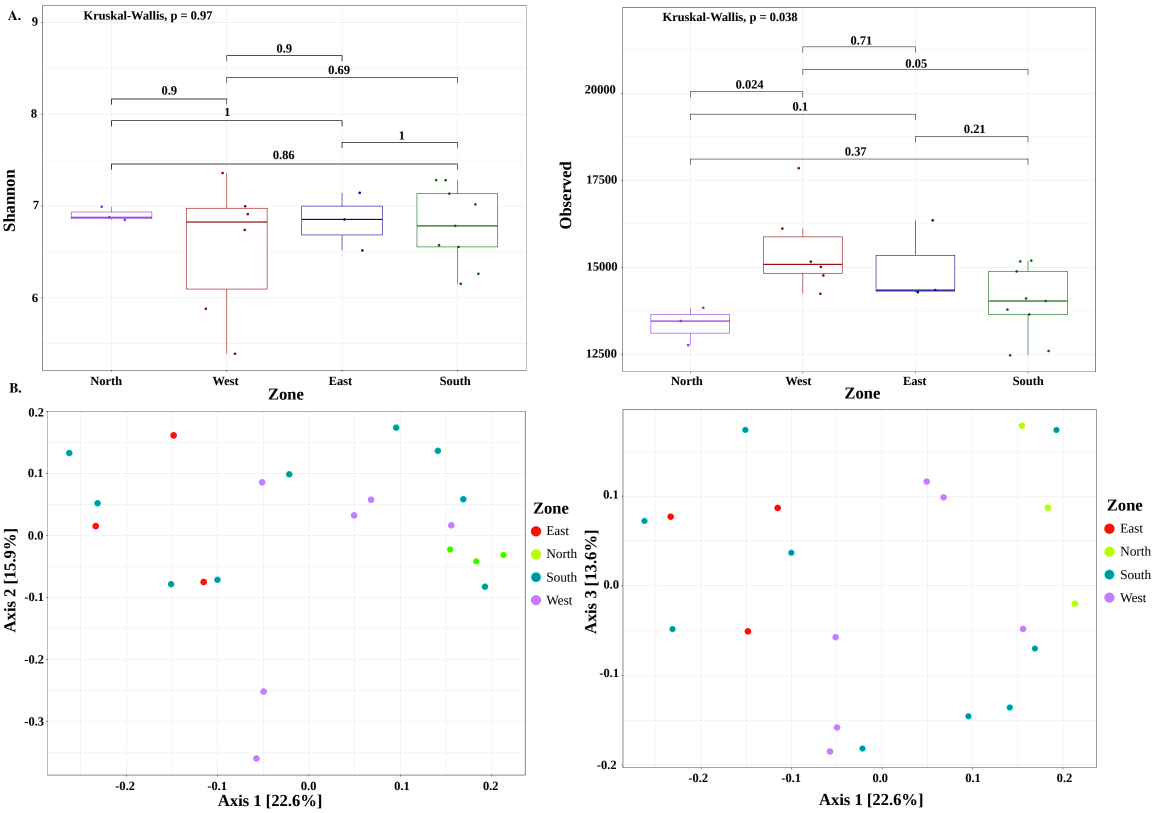


**Fig. S2** **A.** Alpha diversity across zones was measured using the Shannon index and observed species index. The analysis revealed that zones did not significantly influence the microbial diversity of the samples, as anticipated. **B.** PCoA analysis showed partial separation between the West and North zones along Axis 1 and Axis 2. However, when the analysis was performed along Axis 1 and Axis 3, these zones separated more distinctly.

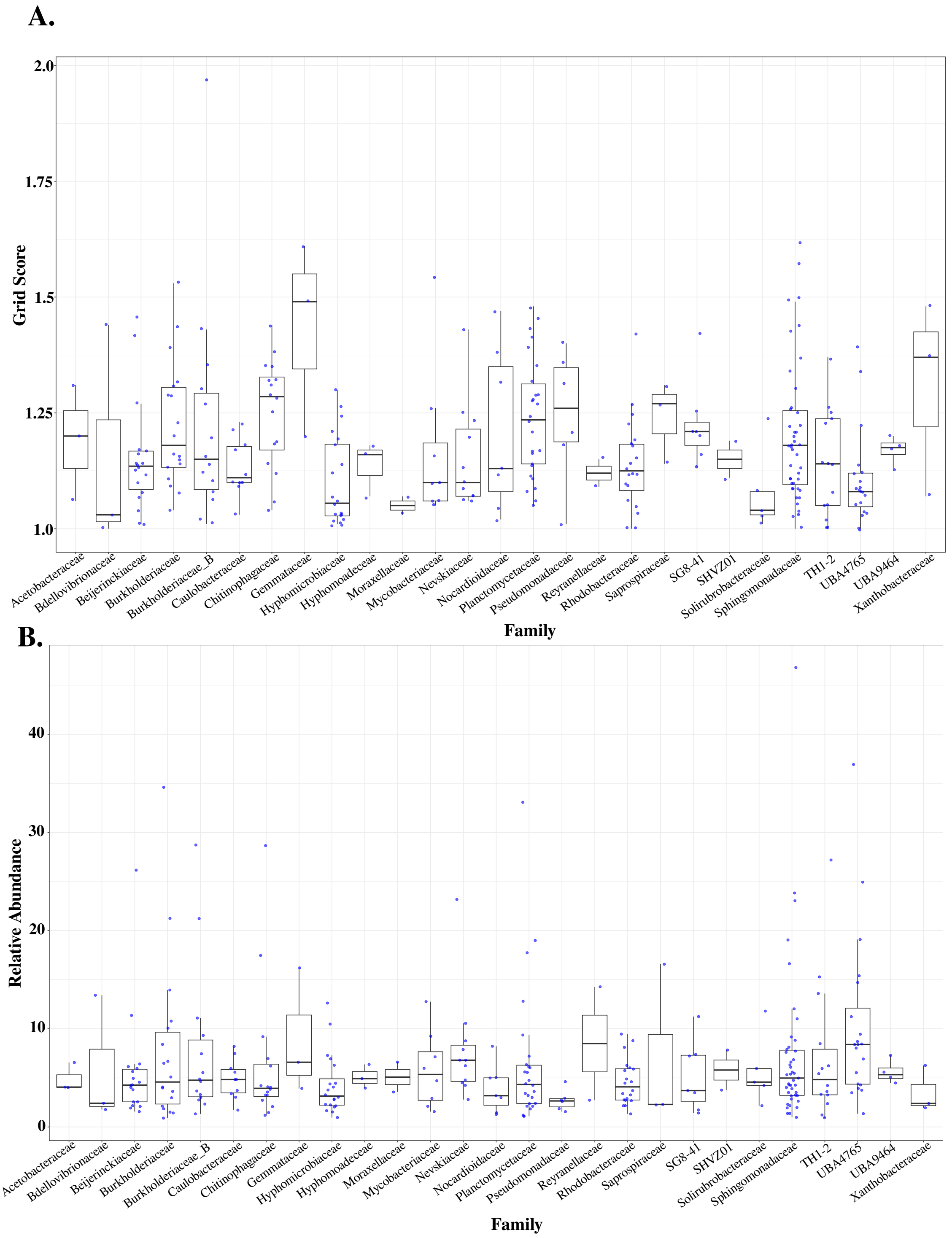


**Fig. S3** Box plot showing the GRiD score **A.** and relative abundance **B.** of Families identified from MAGs. Families with a total relative abundance of less than 10% were excluded from the representation.

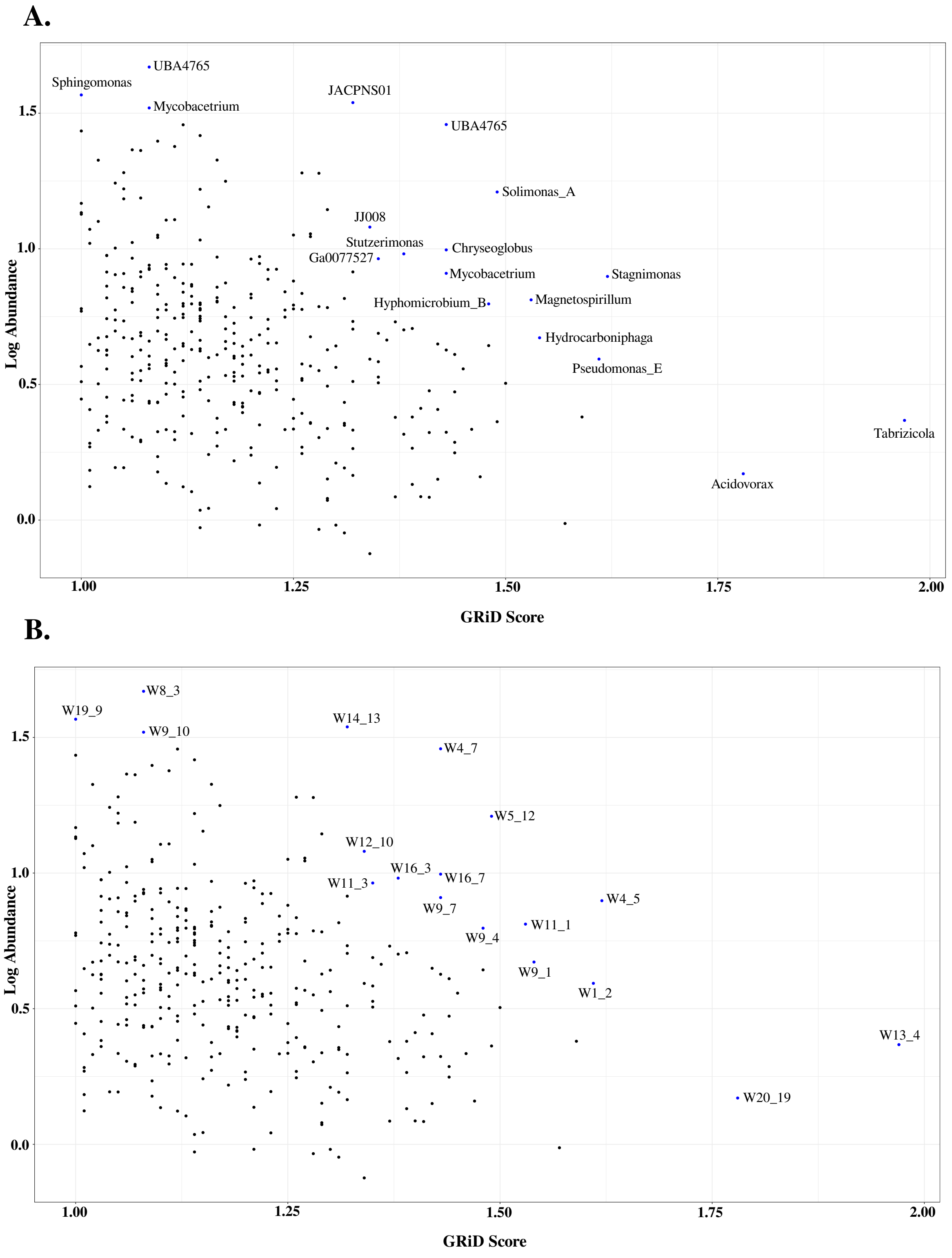


**Fig. S4** Scatter plot illustrating the relationship between GRiD scores and relative abundances of genera **A.** as well as MAG bins **B.** The plot highlights genera with the highest GRiD scores or relative abundances, as well as those with GRiD scores $> 1.25$ and relative abundances $> 0.75$.

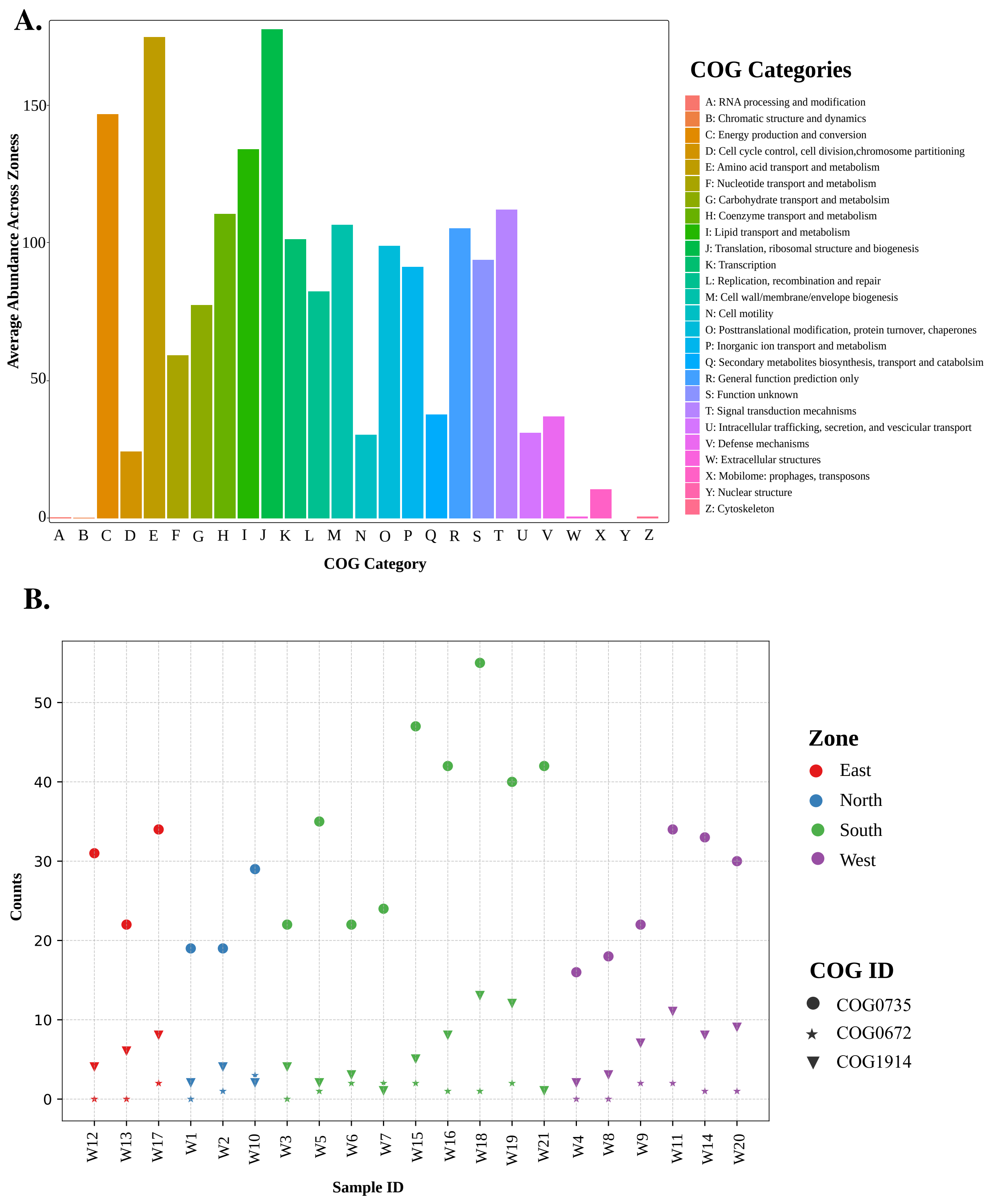


**Fig. S5** **A.** Cluster of orthologous genes in samples: The bar plot represents the average abundance of distinct COG categories across samples. Minimal variation was observed between zones; hence, the average abundance across zones is displayed. **B.** Zone-wise distribution of COGIDs responsible for metal metabolism. COG0735 ($Fe^{2+}$ or $Zn^{2+}$ uptake regulation protein Fur/Zur), COG1914 ($Mn^{2+}$ or $Fe^{2+}$ transporter, NRAMP family), and COG0672 (High-affinity $Fe^{2+}/Pb^{2+}$ permease).

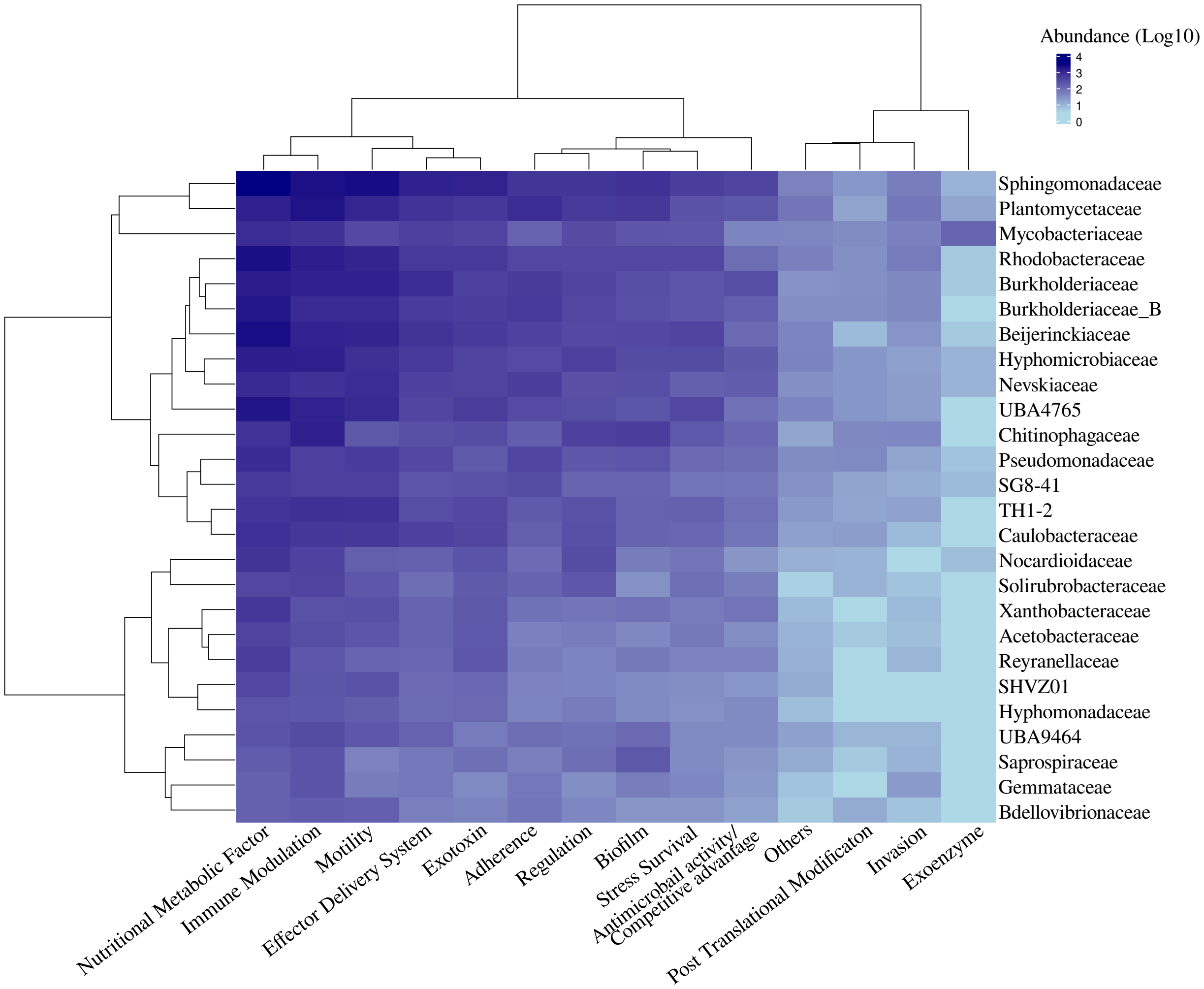


**Fig. S6** Virulence factors distribution in the families identified. Only families with a relative abundance of 10% are shown in this figure. The complete list of virulence genes associated with families is shown in Supplementary Table 2.
